# Pharmacokinetics trumps pharmacodynamics during cocaine choice: a reconciliation with the dopamine hypothesis of addiction

**DOI:** 10.1101/2020.05.20.106096

**Authors:** Ludivine Canchy, Paul Girardeau, Audrey Durand, Caroline Vouillac-Mendoza, Serge H. Ahmed

**Author notes:** Correspondence to: Serge H. Ahmed, Ph.D., Université de Bordeaux, Institut des Maladies Neurodégénératives, UMR CNRS 5293, 146 rue Léo Saignât, 33076 Bordeaux, France.

## Abstract

Cocaine is known to increase brain dopamine at supranormal levels in comparison to alternative nondrug rewards. According to the dopamine hypothesis of addiction, this difference would explain, at least in part, why the latter are eventually given up in favor of continued cocaine use during the transition to addiction. Though resting on solid neuroscientific foundations, this hypothesis has nevertheless proven difficult to reconcile with research on cocaine choice in experimental animals. When facing a choice between an intravenous bolus of cocaine and a nondrug alternative (e.g., sweet water), both delivered immediately after choice, rats do not choose the drug, as would be predicted, but instead develop a strong preference for the nondrug alternative, sometimes to the exclusion of continued drug use. Here we report converging evidence that reconciles this finding with the dopamine hypothesis of addiction. Briefly, our data suggest that cocaine is indeed supranormal in reward magnitude, as postulated by the dopamine hypothesis of addiction, but is less preferred during choice because its pharmacokinetics makes it an inherently more delayed reward than the alternative. Reframing previous drug choice studies in rats as intertemporal choice studies reveals that the discounting effects of delays spare no rewards, including supranormal ones, and that during choice, pharmacokinetics trumps pharmacodynamics. Finally, this study also reveals important gaps in our understanding of drug reward delays that need to be filled by future experimental and theoretical work.

---

Cocaine is known to increase ventral striatal dopamine at supranormal levels in comparison to alternative nondrug rewards, such as, for instance, food or social reward (Di Chiara, 1999; Keiflin & Janak, 2015). According to the dopamine hypothesis of addiction, this initial difference – which persists and even becomes larger with repeated use – would explain, at least in part, why nondrug rewards are eventually given up in favor of continued cocaine use during the transition to addiction (Di Chiara, 1999; Kalivas & Volkow, 2005; Keiflin & Janak, 2015; Redish, 2004; Robinson & Berridge, 2008). Though resting on solid neuroscientific foundations, this hypothesis has nevertheless proven difficult to reconcile with research on drug choices in experimental animals that seems to show the opposite (Ahmed, 2018a; Ahmed, 2018b). When facing a concurrent choice between taking an immediate intravenous bolus of cocaine and having immediate access to a nondrug alternative (e.g., sweet water), rats do not choose the drug, but instead develop a strong preference for the nondrug alternative, sometimes to the complete exclusion of continued drug use (Bagley et al, 2019; Cantin et al, 2010; Kearns et al, 2017; Lenoir et al, 2007; Madsen & Ahmed, 2015; Tunstall & Kearns, 2014; Tunstall & Kearns, 2015; Tunstall & Kearns, 2016; Tunstall et al, 2014). This preference has now been observed with many drugs of abuse, including heroin and methamphetamine (Caprioli et al, 2015a; Caprioli et al, 2015b; Lenoir et al, 2013b; Madsen & Ahmed, 2015; Schwartz et al, 2017; Tunstall et al, 2014; Venniro et al, 2017), with different kinds of nondrug options (Caprioli et al, 2015b; Lenoir et al, 2007; Vandaele et al, 2019; Venniro et al, 2019; Venniro et al, 2018) and also following different drug and/or behavioral histories (Cantin et al, 2010; Caprioli et al, 2015b; Lenoir et al, 2007; Madsen & Ahmed, 2015; Tunstall et al, 2014; Venniro et al, 2018).

Though this discrepancy cries out for an explanation, it has so far been largely overlooked. Recently, we proposed to resolve this discrepancy by considering some of the inherent differences that exist between drug and nondrug options, particularly in reward delays (Ahmed, 2018a; Ahmed, 2018b). Though every effort was made in previous studies to equalize the parameters of choice between drug and nondrug rewards (e.g., same effort; same probability), it has remained particularly difficult to equalize reward delays. Specifically, though the drug and nondrug options are both delivered immediately after a choice, pharmacokinetics prevents the drug to produce its rewarding effects immediately (Lau & Sun, 2002; Pan et al, 1991; Welling, 1986), even when rapidly infused by the intravenous route, as was the case in previous drug choice studies (i.e., typically within less than 5 s). In fact, even if the drug were delivered instantaneously to the venous blood stream, there would remain the incompressible delays of drug distribution, diffusion and action into the brain (Welling, 1986; Wise & Kiyatkin, 2011a). Thus, it is likely that cocaine reward is in fact more delayed than the nondrug reward, though by how much exactly is currently uncertain (see below).

Differences in delays between different options can exert profound influence on choice outcomes in humans and other animals (Ainslie, 1975; Mazur, 1997; Stevens & Stephens, 2010; Vanderveldt et al, 2016). In particular, rats are highly sensitive to reward delays; they typically prefer a small reward now over a larger reward that is delayed by only few seconds (Bradshaw & Szabadi, 1992; Cardinal et al, 2000; Evenden & Ryan, 1996; Logan, 1965; Peterson et al, 2015; Richards et al, 1997; van Gaalen et al, 2006). For instance, when hungry rats are facing a choice between 1 food pellet delivered immediately and 4 food pellets delivered after a delay of 20 s, they typically choose the smaller amount of food, ending up eating less (van Gaalen et al, 2006). Regardless of the mechanisms underlying these delay effects, the pharmacokinetic delays associated with cocaine reward, if sufficiently long, could thus explain rats’ preference for the nondrug alternative in previous drug choice studies, without challenging the dopamine hypothesis of addiction (Ahmed, 2018a; Ahmed, 2018b). Reframing drug choice studies as intertemporal choice studies between a delayed drug reward and an immediate nondrug alternative may reveal that the known discounting effects of delays spare no rewards, including supranormal ones, in rats.

One way to test this delayed drug reward hypothesis could be to try to equalize the delay of the two options by increasing the delay to the nondrug reward, the inverse (i.e., reducing the delay to drug reward to make it quasi-immediate) being not feasible (see also Discussion). For instance, previous research has shown that increasing the delay to the nondrug option can increase drug choice (Lenoir et al, 2007) and eventually shift preference toward the drug (Panlilio et al, 2017; Secci et al, 2016; Venniro et al, 2018). However, without precise knowledge of the delay to drug reward, this approach remains inconclusive because we ignore when the delay added to the nondrug option becomes equal to the unknown delay to drug reward. Thus, knowing the delay to drug reward seems a prerequisite for the validity of this approach. However, as it will soon become apparent, addressing this question is not as simple as it may appear. For comparison, the delay to a nondrug reward is the time interval between a choice response and reward consumption, two observable behavioral events. For instance, with the sweet water option used in our previous choice studies, it is the time interval between the operant response and the onset of sweet water consumption in an adjacent drinking cup (Lenoir et al, 2007). Of particular note, when there is no programmed delay, this time interval is close, but not equal to 0, because it takes some time to move between the response location and the adjacent reward location (i.e., typically within 1-2 s) (Lenoir et al, 2007).

It is not easy to apply this behavioral definition to intravenous cocaine or to any other intravenous drugs of abuse for that matter. Though we know precisely when an intravenous drug infusion begins and ends (and the interval between the two is usually of few seconds), we ignore when exactly drug reward begins (Ahmed, 2018a; Ahmed, 2018b; Aragona, 2011; Wise & Kiyatkin, 2011b). We only know from pharmacokinetics that it cannot start with the drug infusion. After a rat has made a response that triggers an immediate drug infusion, there is no observable behavior that could mark or report unambiguously the onset of the drug reward experience. We previously reported that it takes about 6 s to observe an overt behavioral reaction following the onset of an intravenous cocaine infusion (Lenoir et al, 2007), but we ignore if this reaction precedes, coincides with or even follows the onset of drug reward. As will be explained later, there may be other indirect ways to estimate this delay, but each has its own problems. Thus, the exact delay of onset of cocaine reward remains largely an unknown.

We thus need an approach to test the delayed drug reward hypothesis without prior precise knowledge of the delay to cocaine reward. Previous research has shown that when a sufficiently long delay is added both to a small, immediate reward *and* to a larger, delayed reward, rats change their preference in favor of the larger reward, as if its longer delay no longer mattered (Vanderveldt et al, 2016). For instance, in one paradigmatic experiment, a rat that preferred 2 food pellets immediately over 4 pellets delayed by 20 s reversed its preference to the larger reward when a delay of 15 s was added to the two options (Green & Estle, 2003). Importantly, this change occurred despite the fact that the larger reward was still more delayed by 20 s than the smaller one (i.e., the rat now preferred 4 pellets after 35 s over 2 pellets after 15 s) (Green & Estle, 2003). This preference reversal phenomenon has been observed in pigeons (Ainslie & Herrnstein, 1981; Beeby & White, 2013; Green et al, 1981), in rats (Bradshaw & Szabadi, 1992; Green & Estle, 2003; Ito & Asaki, 1982; Krebs & Anderson, 2012), in monkeys (Kobayashi & Schultz, 2008) and in humans (Green & Myerson, 2004; Kirby & Herrnstein, 1995), and is thought to be a consequence of hyperbolic delay discounting (Ainslie, 1975; Green & Myerson, 2004), though other plausible explanations exist (Blanchard et al, 2013; Sozou, 1998; Stevens & Stephens, 2010).

Regardless of the underlying mechanisms, however, the preference reversal phenomenon suggests a unique way to test the delayed drug reward hypothesis. Thus, if cocaine reward is really larger in magnitude than sweet water reward, but is not chosen because it is more delayed, then adding a sufficiently long programmed delay to these two options should cause rats to shift their choice from sweet water to cocaine, even if the latter remains more delayed. If instead cocaine reward is not larger in magnitude than sweet water reward, then no preference reversal should be observed.

The goal of the present study was to test this unique prediction. We first conducted a systematic analysis of the literature to try to obtain more precise information on the delay of action of intravenous cocaine on ventral striatal dopamine parameters (i.e., dopamine uptake inhibition and extracellular concentrations). Though there is a debate on the exact behavioral significance of these parameters, they are arguably one of the earliest markers of cocaine reward in the brain (Aragona, 2011; Volkow et al, 2019; Wise & Kiyatkin, 2011a). We complemented this research with a behavioral measurement approach. We reasoned that rats should learn about the delay to cocaine reward during cocaine self-administration. Therefore, their response time to a drug omission should be a reflection, albeit an indirect one, of this delay. Finally, we conducted a preference reversal experiment, as outlined above. Overall, our findings confirm the delayed drug reward hypothesis and reconcile previous drug choice studies with the dopamine hypothesis of addiction. There is little doubt that cocaine reward is supranormal in magnitude, but its inherently long delay makes it a less preferred option during choice.

## Materials and Methods

### Systematic analysis of the literature

#### Literature search strategy

We performed a systematic search of the literature for primary research articles that have reported data on the time course of the effects of intravenous cocaine on ventral striatal dopamine parameters with sufficient temporal resolution. To this end, we searched the PubMed Database from inception to January 2020 with the following search expression: “intravenous AND cocaine AND rat AND (onset* OR time-course OR acute OR kinetic* OR pharmacokinetic* OR pharmacodynamic* OR delay*)”. The scope of this initial search was relatively large in an attempt not to miss any potentially relevant article. Finally, the search was limited to studies published in the English language.

#### Selection criteria and additional search

In total, this search strategy allowed us to retrieve a total of 321 articles. Review articles were immediately excluded (n = 11) from the analysis along with one retracted article. The other articles (n = 309) were then reviewed and screened for inclusion or exclusion using the following selection criteria:

1. Cocaine was the drug under study;
2. Rats were the model organism under study;
3. The dependent variable was a ventral striatal dopamine parameter;
4. Cocaine was administered via a jugular vein (as in drug choice studies);
5. Cocaine was administered passively (to avoid potential confounding with conditioned operant response-related dopamine signals);
6. A time course was reported with sufficient temporal resolution (i.e., time intervals between successive measurements < 60 s);

In the end, 305 papers had to be excluded, leaving a total of 4 eligible papers (Figure 1). This initial search was completed with a detailed inspection of the reference lists of these eligible articles in search for additional articles that met the selection criteria. Finally, we also looked for eventual additional items using the tools “Similar articles” and “Cited by” available on the PubMed Database. This additional search allowed us to retrieve 9 additional eligible articles, amounting to a total of 13 articles for final data collection and analysis (Table 1).

**Table 1.** Selected articles after systematic analysis of the literature

| References | # Obs | Infusion (s) | Dose (mg/kg) | DA parameters | NAC region | Methods |
| --- | --- | --- | --- | --- | --- | --- |
| Aragona et al. 2008 | 4 | 6 | 2 | [DA] | Shell & Core | FSCV |
| Cheer et al. 2007 | 1 | 20 | 3 | [DA] | Shell | FSCV |
| Dreyer et al. 2016 | 2 | 6 | 3 | [DA] | Shell & Core | FSCV |
| Espana et al. 2008 | 3 | 2 | 0.75 - 3 | DA UI | Core | FSCV |
| Gratton & Wise 1994 | 1 | 10 | 0.7 - 1 | [DA] | Core | FSCV |
| Heien et al. 2005 | 4 | 20 | 0.3 - 3 | [DA] | Shell | FSCV |
| Mateo et al. 2004 | 2 | 2 | 1.5 | [DA] & DA UI | Core | FSCV |
| Samaha et al. 2004 | 1 | 5 | 2 | DA UI | Core | CV |
| Singer et al. 2017 | 2 | 3 | 1 | [DA] | Shell | FSCV |
| Somers et al. 2009 | 4 | 5 | 3 | [DA] | Shell | FSCV |
| Stuber et al. 2005 | 1 | 6 | 0.3 | [DA] | Core | FSCV |
| Wightman et al. 2007 | 1 | 20 | 1 | [DA] | Shell | FSCV |
| Yorgason et al. 2011 | 1 | 2 | 1.5 | DA UI | Core | FSCV |

Each selected article is referenced in the reference list (Aragona et al, 2008; Cheer et al, 2007; Dreyer et al, 2016; España et al, 2008; Gratton & Wise, 1994; Heien et al, 2005; Mateo et al, 2004; Samaha et al, 2004; Singer et al, 2017; Somers et al, 2009; Stuber et al, 2005; Wightman et al, 2007; Yorgason et al, 2011). The number of observations (# Obs) corresponds to the number of independent measurements reported in the referenced article. The table also shows the range of infusion durations and doses used across the different studies. Dopamine concentration ([DA]) or dopamine uptake inhibition (DA UI) was measured in either the core or shell of the nucleus accumbens using *in vivo* fast-scan cyclic voltammetry (FSCV) or more rarely continuous voltammetry (CV).

**Figure 1:**
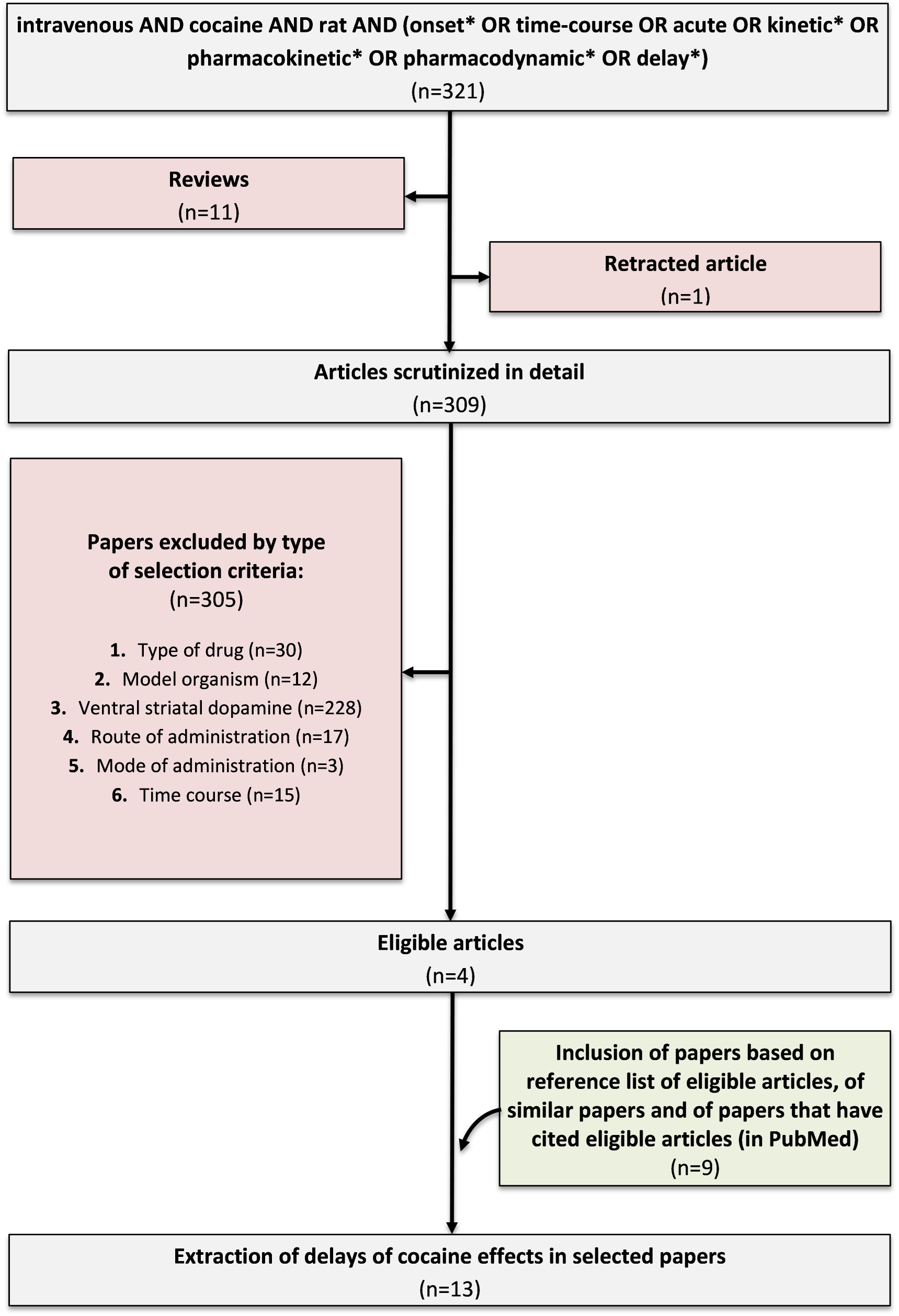
Systematic analysis of the literature. This diagram represents the search and selection process of the primary research articles that were used for extraction of delays of cocaine effects on ventral striatal dopamine. In the end, 13 articles were retrieved that met the selection criteria of this study.

#### Data collection

To quantify the time course of the effects of intravenous cocaine on ventral striatal dopamine parameters, we measured the following delays after onset of drug infusion: 1) delay to reach 10% of the peak effect (D10%); 2) delay to reach 50% of the peak effect (D50%); and 3) delay to reach the peak effect (D100%). D10% was used as the earliest possible proxy of the delay to cocaine reward. We initially tried to obtain these delays directly from the text of the selected research articles but it was only available in 4 articles. In the other 9 articles, we estimated it from the figures using the tool “Measure” of the software Adobe Acrobat Reader DC (Version 2018.011.20055) (see Figure 2A, for an illustration of the method). Briefly, we first measured the length of the time unit (i.e., X-axis) and the height of the unit of the dependent variable (i.e., Y-axis). We then measured the height of the peak effect from baseline (i.e., first highest point of the curve) and computed its 10% and 50% height values. Next, we located on the curve the points corresponding to these height values and draw from them a vertical line to the X-axis to obtain their corresponding length values and thus their delays. In few cases (4 cases), the recordings stopped before a peak effect was reached.

**Figure 2:**
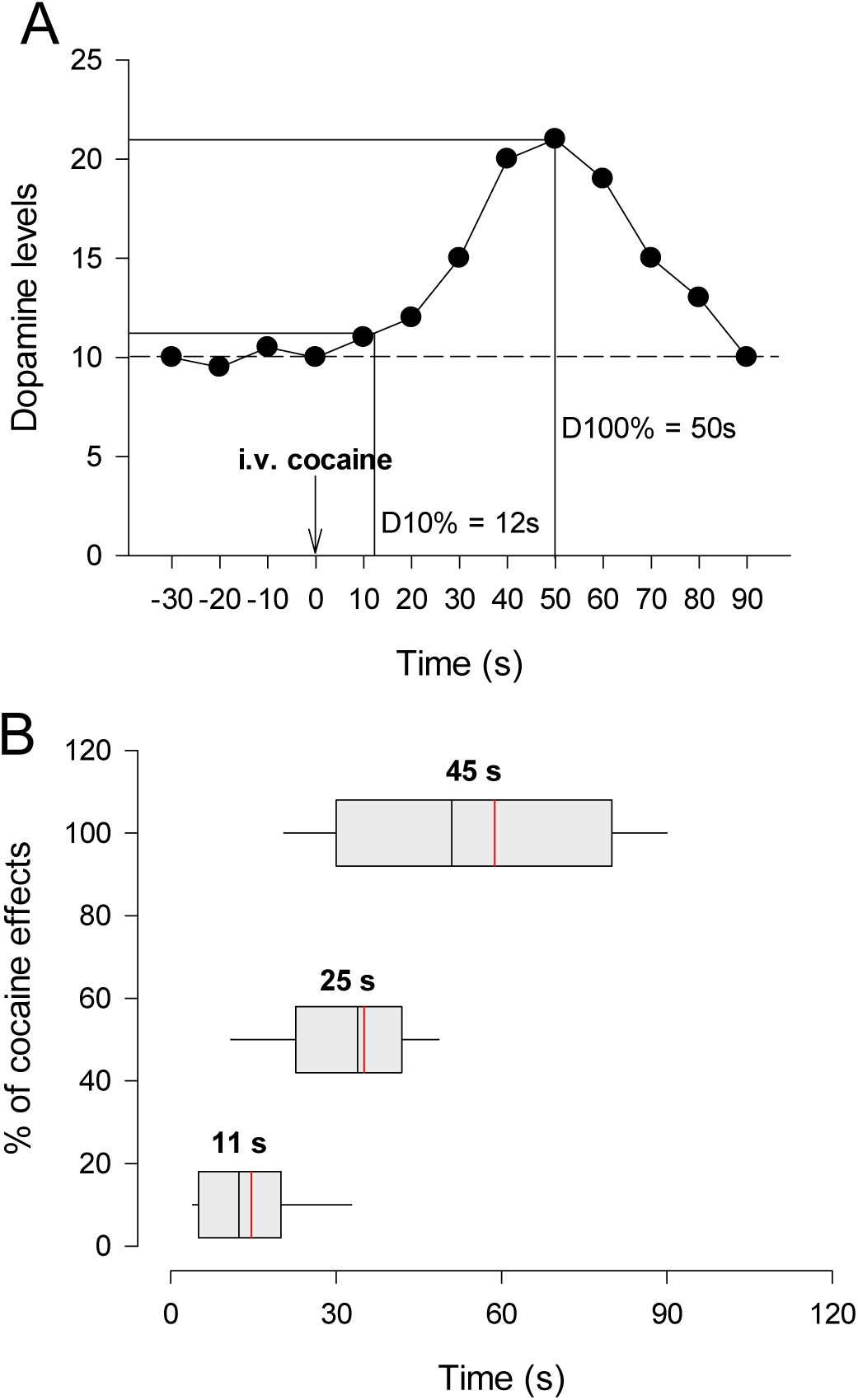
Delays of effects of intravenous cocaine on ventral striatal dopamine. **A**. Schematic illustration showing how D10% and D100% delays were extracted from a figure in a paper selected from the literature. The curve in the figure represents the time course of the effects of cocaine on ventral striatal dopamine levels. First, the units of each axis were converted in distance. On the horizontal time axis, the onset of the intravenous cocaine injection (downward arrow) was used as the origin. A horizontal baseline was then traced from which the height and length of the peak effect (i.e., 100%) were measured. Next, a second horizontal line was traced above the baseline, at a height corresponding to 10% of the peak height. The length of the point of intersection of this 10% line with the curve was then measured. Finally, the lengths of the 10% and 100% effects were converted back in the corresponding time units to obtain the D10% and D100% delays (12 and 50 s, respectively, in the example). A similar method was used to measure the D50% delay (not shown). **B**. Box plots for all estimated delays of effects (i.e., D10%, D50% and D100%). Boxes represent the first, second (i.e., median) and third quartiles; vertical red lines within boxes represent the means; error bars represent the 5 and 95 percentiles. The corresponding statistically indistinguishable shortest delays are indicated above each box.

We thus considered the longest time point on the X-axis as the D100% and the highest point on the Y-axis at the peak effect. Finally, in one study, only the delay to reach 20% of the final effects was reported (Stuber et al, 2005). For convenience, however, this unique case was considered as a D10% measurement.

### Behavioral experiments

#### General experimental procedures

##### Animals and Housing

A total of 83 adult male Wistar rats (225-250 g at the beginning of experiments, Charles River, Lyon, France) were used. Rats were housed in groups of 2 and were maintained in a light-(reverse light-dark cycle), humidity-(60 ± 20%) and temperature-controlled vivarium (21 ± 2°C). All behavioral testing occurred during the dark phase of the light-dark cycle. Food and water were freely available in the home cages throughout the duration of the experiment. Home cages were enriched with a nylon gnawing bone and a cardboard tunnel (Plexx BV, The Netherlands). 28 rats did not complete the behavioral experiments which lasted several months, thereby leaving a total of 55 rats for final analysis. Rats did not complete the experiments due to a variety of factors (e.g., failure to acquire cocaine self-administration; infection; catheter failure).

All experiments were carried out in accordance with institutional and international standards of care and use of laboratory animals [UK Animals (Scientific Procedures) Act, 1986; and associated guidelines; the European Communities Council Directive (2010/63/UE, 22 September 2010) and the French Directives concerning the use of laboratory animals (décret 2013-118, 1 February 2013)]. The animal facility has been approved by the Committee of the Veterinary Services Gironde, agreement number A33-063-922.

##### Surgery

Rats were surgically prepared with an indwelling Silastic catheter in the right jugular vein under deep anesthesia. Behavioral testing commenced at least 7 days after surgery. Additional information about surgery and post-operative care can be found elsewhere (Lenoir et al, 2013a).

##### Apparatus

Fourteen identical operant chambers (30 × 40 × 36 cm) were used for all behavioral testing and training (Imétronic, Pessac, France). Each chamber was equipped with two retractable metal levers on opposite panels of the chamber, and a corresponding white cue light was positioned above each lever. A drinking cup was located 6.5 cm from the left of each retractable lever on the same panel and 6 cm above the grid floor. There were also two syringe pumps, one used to deliver intravenous cocaine via a Tygon infusion line tethered flexibly to the animal’s catheter, the other to deliver sweet water to one of the drinking cup via Silastic tubing. Additional details about apparatus can be found elsewhere (Madsen & Ahmed, 2015).

##### Drugs

Cocaine hydrochloride (Coopération Pharmaceutique Française, Melun, France) was dissolved in 0.9% NaCl, filtered through a syringe filter (0.22 µm) and stored at room temperature (21 ± 2°C). Sucrose (Sigma-Aldrich, St Quentin-Fallavier, France) was dissolved in tap water at room temperature. The sweet solution was renewed each day.

##### Initial cocaine self-administration training

One week after intravenous surgery, rats were first habituated during two 2-h or 3-h daily sessions to the operant chambers. During habituation, no lever or light cue was presented, and rats were allowed to move freely and explore the cage. After habituation, rats were trained to press a lever to self-administer intravenous cocaine during 18-19 daily sessions on a fixed-ratio (FR) 1 time-out 20s schedule of reinforcement, as described elsewhere (Ahmed & Cador, 2006; Ahmed & Koob, 1998; Freese et al, 2018). Specifically, each FR1 response triggered the immediate activation of the cocaine syringe pump which delivered a unit volume of 0.16 ml of a solution containing 0.25-mg cocaine over 4 s. Each daily self-administration session began with extension of one lever and ended after 3 h. The other retractable lever remained retracted. Responding on the lever triggered the immediate activation of the syringe pump that delivered the cocaine bolus and also the light cue above the extended lever for 20 s.

### Specific experimental procedures

#### Response times to drug reward omissions

The rationale behind this test is based on well-established prior knowledge on intravenous cocaine self-administration under a FR1 schedule of reinforcement (Ahmed & Koob, 2005; Pickens et al, 1978; Wise, 1987; Yokel, 1987). Briefly, during acquisition of cocaine self-administration, rats learn that each response is rewarded by cocaine. They then use this information to regulate their intake of cocaine, probably to maintain its rewarding effects around some preferred level and also to avoid overdosing. Typically, after acquisition, they begin a session by responding at a rapid rate to load on cocaine. This initial, brief loading period is then followed by a maintenance period during which rats respond at regular intervals until the end of the session. The length of these regular intervals depends on the unit dose of cocaine available for self-administration: the larger the dose, the longer the inter-response interval. For instance, at the dose used here during training (i.e., 0.25 mg per injection), this interval lasts on average about 280 s in our experimental conditions. Importantly, during the maintenance period, rats rarely respond during the 20-s time-out period that immediately follows the rewarded response, a behavior that is also observed when the time-out period is not signaled.

The relative absence of responding during the post-response time-out period could suggest that after completion of FR1 responding, rats experience quasi-immediately cocaine reward and/or some conditioned interoceptive drug cues. Alternatively and more in accordance with the present hypothesis, it could also indicate that rats have learned to expect that cocaine reward will only occur after a long post-response delay. One way to disentangle these two possibilities is to measure how fast rats respond after drug reward omissions during the maintenance phase of cocaine self-administration. In theory, if responding were followed immediately by cocaine reward and/or some conditioned interoceptive drug cues, then rats should rapidly detect the drug reward omission and thus respond again rapidly (i.e., within few seconds). In contrast, if they have learned that cocaine reward is associated with a long delay, then they should wait a period of time that is at least as long as the drug reward delay before responding again after a drug omission. Since it should take some time to become confident that a drug reward omission has occurred, the omission response time should be slightly longer than the real delay to cocaine reward.

To measure the omission response time, rats were initially trained to self-administer cocaine (0.25 mg per injection) during 18 sessions, as described above. Then, starting from session 19 onward, the time-out cue was removed. This was done to avoid any external influence on responding. As expected, removal of the time-out cue had no significant impact on the maintenance of cocaine self-administration (data not shown). Rats were then tested without the time-out cue during 7 additional sessions of cocaine self-administration before sessions with drug reward omissions began. These sessions were identical to preceding training sessions of cocaine self-administration, except that cocaine reward was occasionally omitted after some FR1 responses during the maintenance period, forcing rats to respond a second time to obtain it. The omission response time was the interval between these two consecutive responses. Drug reward omissions were unsignaled and occurred aperiodically after every 5-7 rewarded responses. No drug reward omissions occurred during the loading period (i.e., first 5 injections). The number of drug reward omissions was initially set to 5 per 3-h session (i.e., first 4 warming-up sessions). Subsequently, to obtain more measurements of omission response times per session, the daily session length was increased to 4.5-h which allowed to test 8 drug reward omissions. Rats were tested under this condition during 10 consecutive sessions, amounting to a maximum of 80 measurements of omission response time per individual rat. Finally, at the end of the experiment, the unit dose of cocaine was halved (i.e., 0.125 mg instead of 0.25 mg) to double the number of injections per session and thus the number of drug reward omissions tested per session, from 8 to 16. Rats were tested with this dose during 4 consecutive sessions which allowed a maximum of 64 measurements of omission response time per individual rat. Since a change of dose affects the amount of, but not the delay to cocaine reward, this test was also used to further validate the drug reward omission procedure.

### Preference reversal experiment

#### Alternate operant training

Rats were first trained to self-administer cocaine during 19 daily sessions, as described above. Starting from session 20 onward, they were also trained on alternate days to respond for sweet water by pressing on a different lever. In total, there were 3 sweet water training sessions alternating with 3 sessions of cocaine self-administration before choice testing (Cantin et al, 2010; Freese et al, 2018; Lenoir et al, 2007; Vandaele et al, 2016). On sweet water sessions, the lever not previously associated with cocaine (i.e., lever S) was extended to mark the onset of the session and to signal sweet water availability; the lever previously associated with cocaine (i.e., lever C) remained retracted. Responding on lever S was rewarded by a 20-s access to water sweetened with 10% sucrose delivered in the adjacent drinking cup on the same wall and was signaled by illumination of the cue-light above lever S for 20 s. Specifically, each FR1 response triggered the immediate activation of the sucrose syringe pump which filled the drinking cup with 0.08 ml of sweet water within 3 s. Additional volumes could be obtained during the remaining 17 s by contacting the cup (i.e., one volume every 2.8 s). The maximum volume that a rat could drink per 20-s access period was 0.32 ml of sweet water. The session ended after rats had earned a maximum of 30 sweet water rewards which typically occurred in less than 30 min.

#### Discrete-trials choice procedure

After acquisition of lever pressing for cocaine and sweet water, rats were allowed to choose during several consecutive daily sessions between lever C and lever S on a discrete-trials choice procedure. Each daily choice session consisted of 16 discrete trials, spaced by 10 min, and divided into two successive phases, sampling (4 trials) and choice (12 trials). During sampling, each trial began with the presentation of one single lever in this alternate order: C – S – C – S. Lever C was presented first to prevent an eventual drug-induced taste aversion conditioning or negative affective contrast effects (Lenoir et al, 2007). If rats responded on the available lever within 5 min, this triggered the immediate retraction of the lever, the immediate activation of the relevant syringe pump (i.e., sucrose or cocaine as described above) and the immediate illumination of the cue-light above the sampled lever for 20-s. If rats failed to respond within 5 min, the lever retracted until the next trial. Thus, during sampling, rats were allowed to evaluate each option separately before making their choice. Choice trials were identical to sampling trials, except that they began with the presentation of both levers S and C and ended with their simultaneous retraction. Rats had to respond on one of the two levers to make their choice and obtain the corresponding reward. During sampling and choice, the response requirement was set to 2 consecutive responses to avoid eventual accidental choice. A response on the alternate lever before satisfaction of the response requirement reset it. Response resetting occurred very rarely, however. Rats were tested in this discrete-trials choice procedure during 12 daily sessions before being tested with different programmed delays.

#### Adding a programmed delay to both options

After stabilization of preference, the same programmed delay was added to both options using a choice procedure identical to that described above. Thus, when rats responded during sampling or choice, this triggered the immediate retraction of the lever(s) and the immediate illumination of the relevant cue-light for 20 s (+ programmed delay), but the relevant syringe pump (i.e., cocaine or sucrose) was activated only after a programmed delay. Four programmed delays were tested in the following ascending order: 10, 20, 40 and 60 s. Each delay was tested during at least 6 consecutive daily sessions until stabilization of group-average behavior (i.e., no increasing or decreasing trend across 3 consecutive sessions). In total, rats were tested during 35 consecutive sessions. The last 3 baseline choice sessions preceding testing with programmed delays served as the 0-s delay control condition.

#### Data analysis

All data were subjected to relevant repeated measures ANOVAs, followed by Tukey post hoc tests where relevant. Comparisons with a fixed theoretical level (e.g., 50%) were conducted using one sample t-tests. These comparisons were also used to identify for each measured delay the shortest delay that was statistically indistinguishable using a 1-s resolution. Alpha level for detecting statistical significant differences was set to p < 0.05. Statistical analyses were run using Statistica, version 7.1 (Statsoft Inc., Maisons-Alfort, France).

## Results

### Systematic analysis of the literature

A total of 13 studies ended up meeting the criteria of selection (see Table 1). Not surprisingly, all the selected studies measured the effects of intravenous cocaine on ventral striatal dopamine parameters using *in vivo* voltammetry methods which have the required temporal resolution. All recordings were either made in the core or shell of the nucleus accumbens. Among these studies, some reported several independent measurements of some delays but not of the others (e.g., in one study, there were 3 independent measurements of the D10% and D100% delays, but 0 measurement of the D50% delay). In the end, we were able to extract the following number of measurements per delay across all studies: 27 for the D10% delay, 20 for the D50% delay and 26 for the D100% delay. Missing values were due to a lack of information in the original studies. Overall, there was a relatively large heterogeneity in outcomes across the different studies and observations (Figure 2B).

Regardless of the delays considered, coefficients of variation (CV) were relatively high (i.e., CVs > 60%), attaining 75% for the D10% delay which represents the earliest neurochemical proxy for the delay to cocaine reward. This heterogeneity was not related to the variation in the dose (R2 = 0.027, NS) or in the duration of infusion (R2 = 0.106, NS) used across the different studies (see Table 1). There was also no significant difference as a function of the nucleus accumbens subregion (i.e., core versus shell) (F1,25 = 0.057, NS). Thus, though the measurement methods in the selected studies were homogeneous, the heterogeneity in extracted data precludes any reliable inference of a definite delay to cocaine reward. Despite this limitation, we can nevertheless conclude that this delay is far from being immediate. The mean D10% delay was equal to 14.6 ± 10.3 s (mean ± 1 standard deviation). At the other extreme, the mean D100% was equal to 58.7 ± 36.0 s. Between these two extremes, the mean D50% delay was equal to 35.0 ± 21.8 s. All these delays were significantly different from each other (F2,38 = 41.37, p < 0.001, post-hoc Tukey HSD p-values < 0.001). The shortest delays that were statistically indistinguishable to the D10%, D50% and D100% delays were 11 s [t(26) = 1.82, p > 0.05], 25 s [t(19) = 2.05, p > 0.05] and 45 s [t(25) = 1.94, p > 0.05], respectively (Figure 2B).

### Omission response times during cocaine self-administration

We analyzed for each individual rat the distribution of omission response times as a function of the cocaine dose available (i.e., a maximum of 80 and 64 measured times at 0.25 and 0.125 mg, respectively) (Figure 3A). Overall, for all individuals, most omission response times were relatively long and variable (CVs > 50 %), but they did not change significantly with the dose available (p-values > 0.05). Individual mean omission response times ranged from 24.0 to 88.1 s at the dose of 0.25 mg and from 22.6 to 113.2 at the dose of 0.125 mg. When all individual mean data were pooled, the mean omission response time of the group was 44.5 ± 16.1 s (mean ± 1 standard deviation) for the dose of 0.25 mg and 43.5 ± 22.8 s for the dose of 0.125 mg (Figure 3B). This difference was not statistically significant (F1,16 = 1.89, NS). The shortest delays that were statistically indistinguishable to these two measured delays were 37 s [t(16) = 1.91, p > 0.05] and 32 s [t(16) = 2.08, p > 0.05], respectively. Overall these omission response times are within the interval range defined by the delays D10% and D100% of the effects of intravenous cocaine on ventral striatal dopamine parameters (Figure 2B). For comparison, the mean inter-response interval during the 3 baseline sessions of cocaine self-administration preceding sessions of drug reward omissions was about 10 times longer: 277.4 ± 114.4 s (mean ± 1 standard deviation).

**Figure 3:**
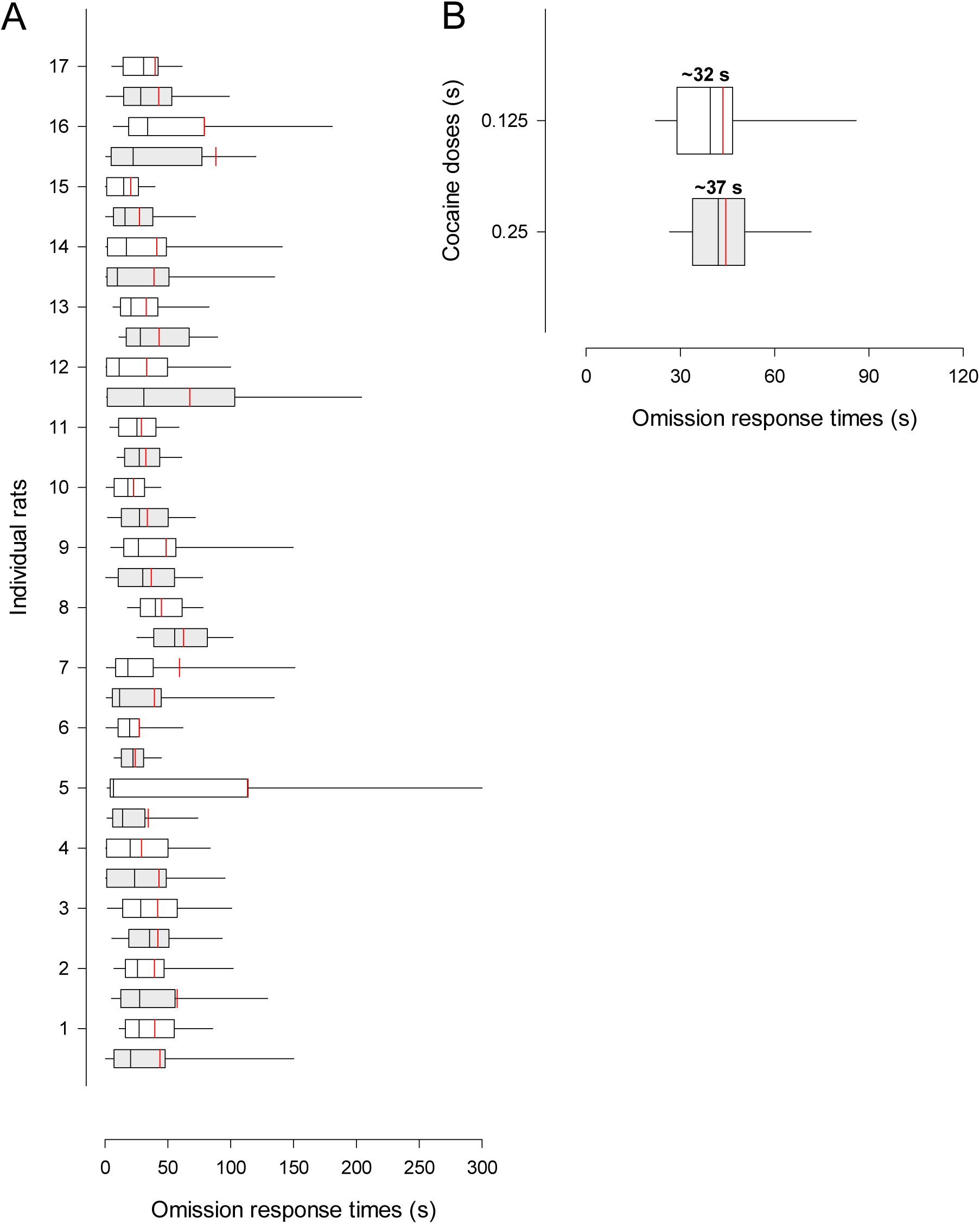
Omission response times during intravenous cocaine self-administration. **A**. Box plots of omission response times for each individual (n = 17) at doses 0.25 mg (light grey) and 0.125 mg (white). Boxes represent the first, second (i.e., median) and third quartiles; vertical red lines within boxes represent the means; error bars represent the 5 and 95 percentiles. **B**. Box plots of all individual mean omission response times at doses 0.25 mg (light grey) and 0.125 mg (white). The corresponding statistically indistinguishable shortest delays are indicated above each box.

If the omission response time is a genuine reflection of the delay to cocaine reward, as hypothesized here, then it should increase after adding a programmed delay between the response and the drug infusion during cocaine self-administration. To test this prediction, we measured in a separate group of rats (n = 16) how different programmed delays influenced omission response times (6 drug reward omissions per 3.5-h session). Three programmed response-infusion delays were tested (0, 20 and 10 s, in that order), each during at least 7 consecutive sessions. Only the results obtained during the last 5 sessions of each condition were considered for analysis. Rats used in this study were initially trained to self-administer cocaine (0.25 mg per injection over 2 s) using a modified FR1 schedule. Briefly, immediately after each response, the lever was retracted for 30 s. Since most rats did not respond before that time after a drug omission, as shown above, this modification should be of little consequence on omission response times. This was indeed the case. When no programmed delay was added between the response and the drug infusion, individual mean omission response times were comparable to those measured in the previous experiment and ranged from 34.9 to 106.9 s (Figure 4A). In addition, the mean omission response time of the group was 59.0 ± 16.9 s which was also comparable to that measured previously and much shorter than the interval between rewarded responses from the same sessions (i.e., 271.0 ± 53.9 s). More importantly, adding a programmed delay between the response and the drug infusion increased the omission response time by a duration that was close, though not identical, to the added delay (i.e., + 14.5 s and + 18.3 s for the 10-s and 20-s programmed delays, respectively) (F2,30 = 8.59, p < 0.01, post-hoc Tukey HSD p-values < 0.01) (Figure 4B). This latter result suggests that when no programmed delay is present, the length of the omission response time should be close to the underlying delay to cocaine reward. Finally and as expected, adding a programmed delay did not affect the mean intervals between rewarded responses during testing (F2,30 = 0.13, NS).

**Figure 4:**
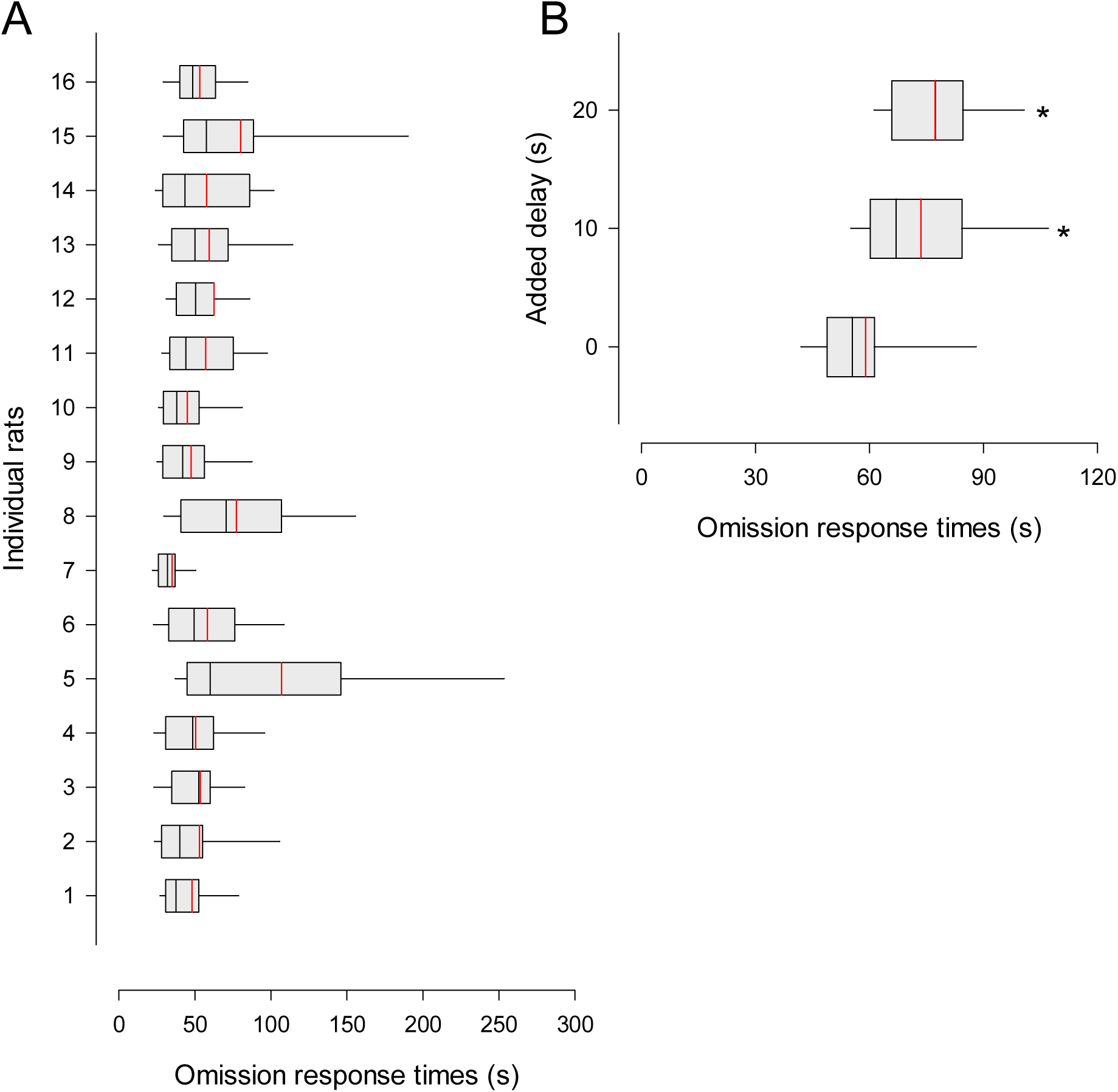
Effects of adding a programmed drug delay on omission response times. **A**. Box plots of omission response times for each individual (n = 16) at the control 0-s delay. Boxes represent the first, second (i.e., median) and third quartiles; vertical red lines within boxes represent the means; error bars represent the 5 and 95 percentiles. **B**. Box plots of all individual mean omission response times as a function of the added delay. * different from added 0-s delay, post-hoc Tukey p-value < 0.05.

### Preference reversal experiment

As expected, when offered a choice between cocaine and sweet water, rats rapidly developed a strong preference for the latter (F11,231 = 7.91, p < 0.01). This preference was manifest from session 2 onward (different from the indifference level of 50%: p < 0.05, t-test) (Figure 5A). On the final 3 sessions, the large majority of individual rats showed either a marked preference for sweet water (n = 19, % cocaine choice < 40%) or were indifferent (n = 3, % cocaine choice between 40-60%) (Figure 5B). No individual rat showed a preference for cocaine in this study (i.e., % cocaine choice > 60%). In addition, all rats made their choice relatively rapidly (i.e., 16.5 ± 6.1 s), indicating little hesitation. Finally, the amount of sweet reward consumed per sweet water choice was maximal for all individuals (i.e., 0.32 ± 0.00 ml).

**Figure 5:**
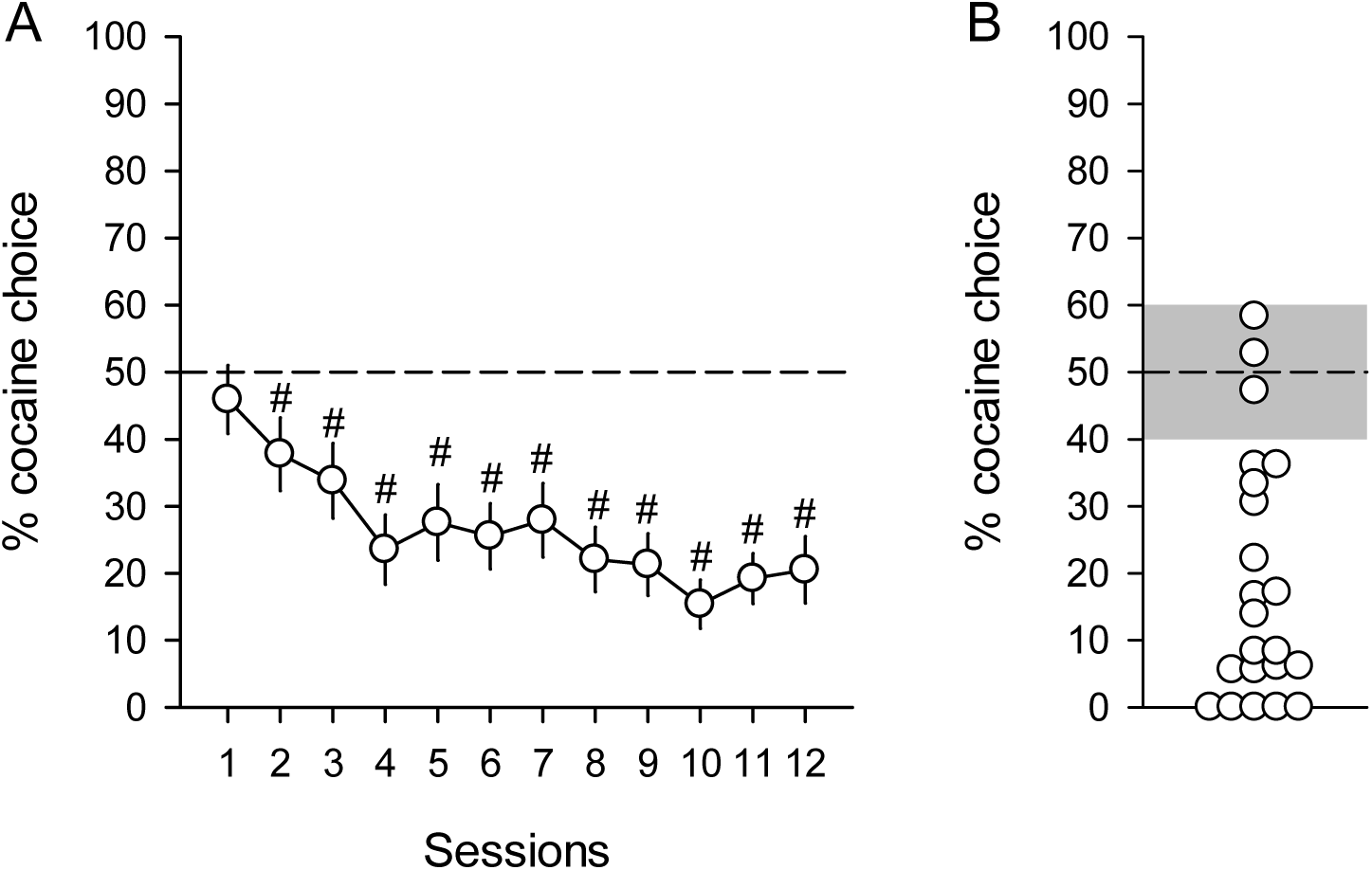
Preference for sucrose over cocaine during choice. **A**. Mean (± s.e.m.) percentage of cocaine choice across sessions. The horizontal dotted line represents indifference level. # different from indifference, t-test p-values < 0.05. **B**. Average individual preferences over the last 3 choice sessions. Each point represents a different individual (n = 22). The grey zone represents ± 10% around the indifference level.

When a programmed delay was added to both options, rats became slower to make their choice and sometimes did not make a choice within the 5-min imparted time (F4,84 = 16.76, p < 0.01) (Figure 6A). As the result, they completed less choice trials (F4,84 = 12.84, p < 0.01) (Figure 6B). This outcome was expectable because adding a programmed delay to the two options should decrease both their values, thereby decreasing the choice stake. In 3 rats, the % of completed trials dropped below 25% when the delay was 40 s or higher, precluding any reliable analysis of their choice behavior at these programmed delays (Figure 6C). These rats were thus excluded, leaving a total of 19 rats for final analysis. Overall, increasing the length of the programmed delay caused rats to increase their choice of cocaine and eventually to reverse their preference for sweet water to cocaine (F4,72 = 39.16, p < 0.01) (Figure 7A). Preference for cocaine tended to level off above 40 s and reached statistical significance at the longest delay of 60 s (above the indifference level of 50%: p < 0.05, t-test). Importantly, this effect was not incidental to a reduction in the amount of sweet reward consumed per sweet water choice (F4,72 = 2.24, NS) which remained close to the maximum available even at the longest delay of 60 s (i.e., 0.30 ± 0.01 ml).

**Figure 6:**
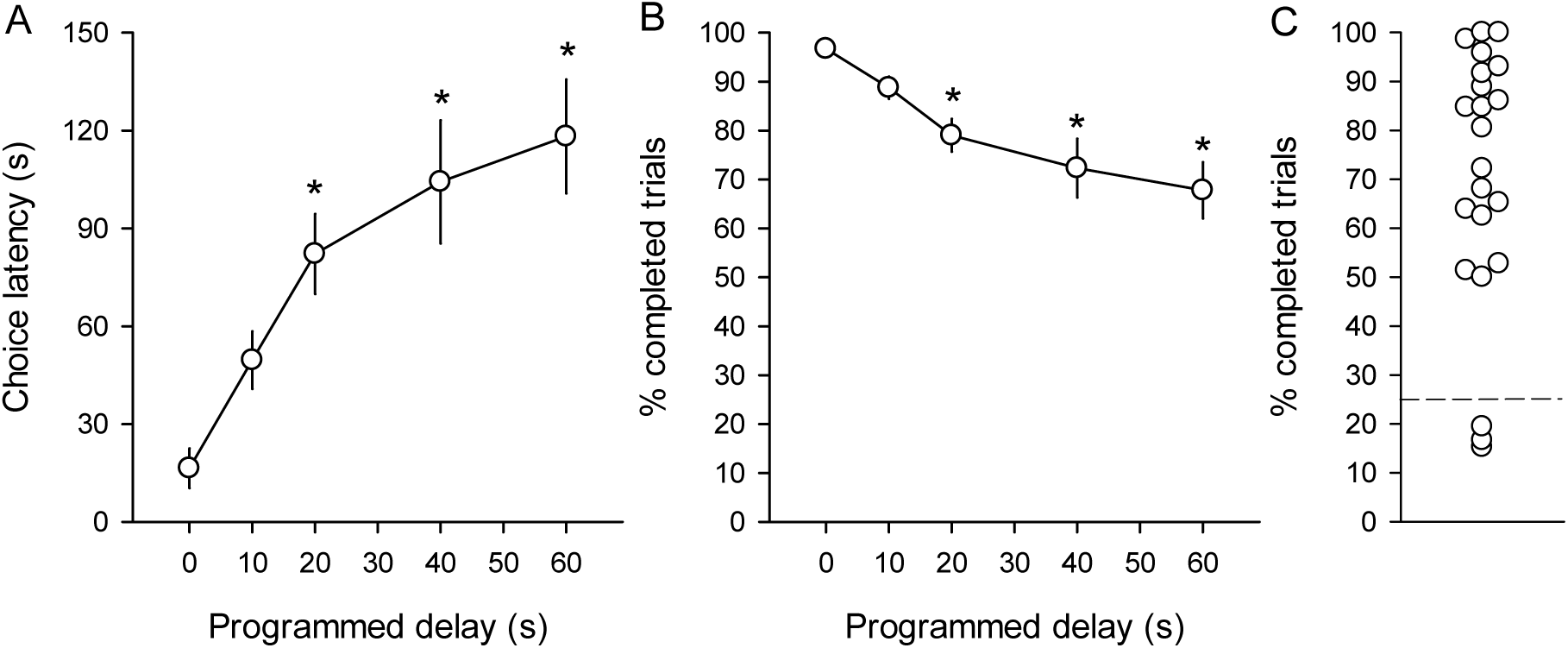
Effects of adding an increasing programmed delay to both options on choice performance. **A**. Mean (± s.e.m.) choice latencies as a function of programmed delays. **B**. Mean (± s.e.m.) % of completed choice trials as a function of programmed delays. **C**. Average % completed trials at the two longest programmed delays. Each point represents a different individual (n = 22). Points below the dotted line represent individuals (n = 3) that completed less than 25% of their choice trials. * different from 0-s delay, post-hoc Tukey p-value < 0.05.

**Figure 7:**
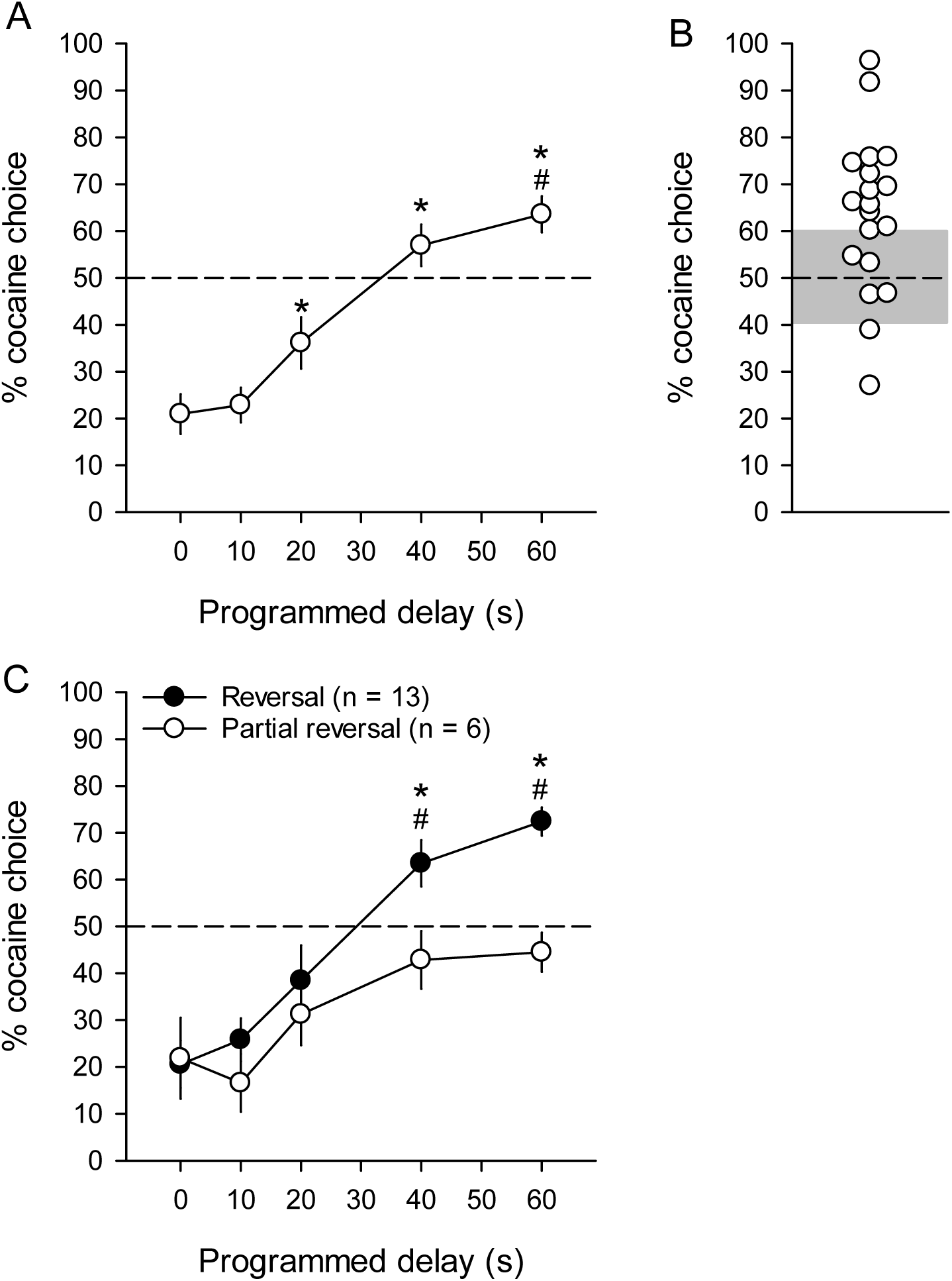
Effects of adding an increasing programmed delay to both options on preference. **A**. Mean (± s.e.m.) % cocaine choices as a function of programmed delays. **B**. Average individual preferences over the last 3 choice sessions with the longest programmed delay of 60 s. Each point represents a different individual (n = 19). The grey zone represents ± 10% around the indifference level. **C**. Mean (± s.e.m.) % cocaine choices in the two subgroups of rats as a function of programmed delays. # above indifference, t-test p-values < 0.05; * different from the other subgroup, post-hoc Tukey p-value < 0.05.

Interestingly, response latencies during sampling trials were generally congruent with and even predictive of subsequent choice behavior. Consistent with previous research (Lenoir et al, 2007), when no programmed delay was added (i.e., during baseline preference for sweet water), rats responded faster for sweet water, the preferred option, than for cocaine (2.79 ± 0.74 versus 9.32 ± 1.52; F1,18 = 17.63, p < 0.01). When a programmed delay was added to both rewards, rats became slower to respond for either reward, but the increase in response latency was steeper for sweet water than for cocaine (F4,72 = 7.72, p < 0.01). At the longest programmed delay, rats now responded faster for cocaine, the now preferred reward, than for sweet water (73.52 ± 19.70 versus 123.36 ± 24.55; F1,18 = 8.30, p < 0.01). Once again, this effect was not incidental to a reduction in the amount of sweet reward consumed per sweet water sampling (F4,72 = 1.98, NS) which remained close to the maximum available even at the longest delay of 60 s (i.e., 0.30 ± 0.01 ml).

Though all rats increased their cocaine choice with increased programmed delays, 13 rats reversed their preference to cocaine at 60 s (i.e., % cocaine choice > 60%) while the other 6 rats either became indifferent (n = 4) or continued to slightly prefer sweet water (n = 2) (Figure 7B). Importantly, when we looked at each of these subgroups separately, we found that the majority of rats that reversed their preference to cocaine continued to increase their cocaine choices with increased delays while the other rats did not (Subgroup: F4,68 = 3.37, p < 0.05). In the latter rats, cocaine choices tended instead to level off around the indifference level at 40 s. It is thus unlikely that they would have reversed their preference to cocaine at longer programmed delays (Figure 7C). The levelling-off of their cocaine choices around indifference may suggest that for these few rats, cocaine is not larger, but similar in reward magnitude than sweet water. There were no other differences between these two subgroups during choice testing. They initially acquired the same preference for sweet water and at the same rate than the majority of rats that subsequently reversed their preference to cocaine (Subgroup: F1,17 = 0.01, NS; Subgroup × Delay: F4,68 = 0.50, NS). In addition, regardless of the programmed delays, rats that did not fully reverse their preference completed as many choice trials as rats that fully reversed their preference (Subgroup: F1,17 = 0.84; Subgroup × Delay: F4,68 = 2.40, NS). They were also equally fast to make their choice (Subgroup: F1,17 = 0.94, NS; Subgroup × Delay: F4,68 = 2.34, NS) and consumed the same amount of sweet reward when they happened to choose this reward (Subgroup: F1,17 = 0.79, NS; Subgroup × Delay: F4,68 = 0.64, NS).

## Discussion

As expected, the systematic analysis of the literature on the delays of action of intravenous cocaine on ventral striatal dopamine parameters has revealed that these delays are relatively long, probably of the order of tens of seconds, as was suggested previously (Wise & Kiyatkin, 2011b). However, the analyzed data were relatively heterogeneous across studies which prevented us from inferring a specific delay. We could not relate this heterogeneity to nucleus accumbens subdivisions (i.e., core versus shell) or to variation in cocaine doses and infusion durations across studies. The latter finding was expectable from known cocaine pharmacokinetics (Pan et al, 1991), at least within the range of infusion durations used in the analyzed studies (see Table 1). We thus tried to complement this research with a behavioral approach which used rats’ response time to drug reward omissions as a reflection of the delay to cocaine reward. Overall, this approach confirmed that the delay to cocaine reward is likely tens of seconds long, though once again data heterogeneity, both between-and within-subject, precluded any inference of a definite delay. Finally, having obtained sufficient converging evidence that cocaine is likely a long delayed reward, we conducted a preference reversal experiment in rats that were choosing between cocaine and sweet water. Adding a programmed delay to these two options caused all rats to increase their choice of cocaine and a large majority of them to shift their preference from sweet water to cocaine. Taken together, this research confirms the delayed drug reward hypothesis (Ahmed, 2018a; Ahmed, 2018b) and offers a reconciliation of previous cocaine choice studies in rats and the dopamine hypothesis of addiction (Di Chiara, 1999; Kalivas & Volkow, 2005; Keiflin & Janak, 2015; Redish, 2004; Robinson & Berridge, 2008). Thus, for most rats, cocaine reward is supranormal in magnitude, but its inherently longer delay makes it a less desirable option during choice in comparison to smaller, sooner nondrug rewards. Since the delay to cocaine reward is imposed by pharmacokinetics, this also suggests that during choice, pharmacokinetics can trump pharmacodynamics, a conclusion that should be taken more into account in future research, notably when comparing drug versus nondrug rewards.

Thus, previous drug choice studies in rats were likely intertemporal choice studies between a delayed drug reward and an immediate nondrug alternative. A major difference with more classic intertemporal choice studies, however, is that the delay to cocaine reward is less accessible and modifiable than the delay of a typical larger, longer nondrug reward is. As explained above, we have evidence that the delay to cocaine reward is relatively long, but we ignore its exact length, mainly because our available data are indirect and relatively heterogeneous across studies and individuals. We can prolong the delay to cocaine reward by retarding a drug infusion, but we cannot shorten it, at least not sufficiently to make it as immediate as a nondrug reward alternative can be. For instance, in a previous study, we reduced the duration of intravenous cocaine infusion from 4 s (which is the duration of infusion used in our drug choice studies, including the present study) to 1 s, but this had no observable impact on drug choice outcomes (unpublished findings). This negative finding was not unexpected, however, since the duration of infusion should have little effect on the time course of drug concentration in the body and the brain, and thus on the delay to cocaine reward, except, of course, when the infusion duration is excessively long (Pan et al, 1991). Thus, it is not possible to offer rats a drug reward with either a known delay or no delay at all, a limitation that considerably restricts our ability to compare drug rewards with nondrug rewards. This analysis identifies an unforeseen gap in our knowledge on drug reward that needs to be filled by new research.

According to the delayed drug reward hypothesis, if one were able to offer cocaine reward without a delay, all rats should choose cocaine and this regardless of the magnitude of the immediate nondrug option available. One way to resolve this problem could be to train rats to self-administer cocaine directly into the ventral striatum via chronic intracerebral cannula (Goeders & Smith, 1987). However, there is little certitude that the rewarding effects of intrastriatal cocaine will reproduce appropriately the rewarding effects of intravenous cocaine which also increases dopamine levels in several other reward-related brain regions. This factor may explain, at least in part, why intra-striatal cocaine was found in previous research to support only weakly self-administration in rats (Carlezon et al, 1995; Goeders & Smith, 1983; Rodd-Henricks et al, 2002). Thus, if rats were offered a choice between intravenous cocaine and intrastriatal cocaine, it is not sure that they would prefer the intracranial route. This analysis nevertheless raises a number of theoretical issues worth pursuing in future research.

There is, however, an alternative approach that may reproduce the rewarding effects of intravenous cocaine reward perhaps more accurately and equally rapidly than direct intracerebral infusions of cocaine. It consists of using optogenetics to photostimulate selectively and quasi-immediately midbrain dopamine neurons (Pascoli et al, 2018; Pascoli et al, 2015), thereby increasing dopamine levels in several brain regions, like intravenous cocaine does but more rapidly. Using this approach, it has recently been found that photostimulation of midbrain dopamine neurons is about 4 times more addictive than intravenous cocaine in mice (Pascoli et al, 2018; Pascoli et al, 2015). Specifically, a large majority of mice (i.e., 70%) developed addition-like behavior to photostimulation of midbrain dopamine neurons while only a minority (i.e., less than 20%) developed this behavior to intravenous cocaine (Pascoli et al, 2015). In addition, when mice were offered a concurrent choice between photostimulation of midbrain dopamine neurons and a normally highly rewarding palatable drink, virtually all of them chose the dopamine neuron stimulation nearly exclusively (C. Lüscher, personal communication). Thus, when the effects of cocaine on brain dopamine levels are reproduced but without the long delays, animals show a strong preference toward them, as predicted by the delayed drug reward hypothesis (Ahmed, 2018a; Ahmed, 2018b). This interpretation also predicts that if given a choice, mice should choose photostimulation of midbrain dopamine neurons over intravenous cocaine – a prediction that remains to be tested. Importantly, this interpretation does not exclude other possible explanations. For instance, the lower addictive effects of cocaine in comparison to photostimulation of midbrain dopamine neurons may also be due to its action on other neurotransmitters in the brain that could mitigate its action on dopamine (Lüscher et al, 2020).

As explained in the Introduction, the delayed drug reward hypothesis was initially developed to explain why rats prefer nondrug reward over intravenous drug reward and reconcile this finding with the dopamine hypothesis of addiction. It is not clear, however, how this hypothesis can also explain the behavior of cocaine-preferring rats that are sometimes observed, though not always (e.g., present study), in drug choice studies (Ahmed, 2010; Ahmed et al, 2013). For instance, when we pooled all our cocaine choice experiments, we found that the prevalence of cocaine-preferring rats was about 10% (Cantin et al, 2010). This prevalence did not increase significantly after extended access to and escalation of cocaine self-administration (Cantin et al, 2010), suggesting that cocaine preference may be a preexisting individual trait. This interpretation is consistent with subsequent research on the neuronal correlates that predict individual drug preferences in rats (Guillem & Ahmed, 2018). What distinguishes drug-preferring rats from the majority of nondrug-preferring ones remains currently poorly understood, however. We previously suggested that these individuals may be homologous to the minority of cocaine users who go on to develop addiction in humans (Ahmed, 2010; Ahmed et al, 2013). However, the delayed drug reward hypothesis suggests another, more counterintuitive possibility. Cocaine-preferring rats could simply be less sensitive to long reward delays compared to the majority of rats. If true, they should also behave less impulsively in other intertemporal choice settings, choosing more often the larger, more delayed options than the other rats. In other words, low choice impulsivity would predispose to greater drug choice in rats. Obviously, this is the opposite of what we know about the relationship between choice impulsivity and vulnerability to drug use and addiction in people (Bickel et al, 2014; de Wit, 2009; Rung et al, 2019; Volkow & Baler, 2015). People who behave less impulsively in intertemporal choice settings are less, not more, vulnerable to use drugs and go on to develop addiction. Clearly, further research is needed to resolve this apparent translational gap between rats and humans [for additional discussion of this translational gap, see (Ahmed, 2018b; Ahmed, 2019)].

In conclusion, the present study provides strong evidence for the delayed drug reward hypothesis and offers a reconciliation of cocaine choice studies in rats and the dopamine hypothesis of addiction. Thus, for most rats, cocaine reward is supranormal in magnitude, but its inherently long delay makes it a less preferred desirable option during choice in comparison to nondrug rewards. Since the delay to cocaine reward is imposed by pharmacokinetics, this suggests that pharmacokinetics can trump pharmacodynamics in drug self-administration studies, a conclusion that should be taken more into account in future research, notably when comparing drug versus nondrug rewards. Finally, this study also reveals several important knowledge gaps, notably in our understanding of drug reward delays, that need to be filled by future experimental and theoretical research.

## Acknowledgements

We thank Christophe Bernard, Mathieu Louvet and Eric Wattelet for administrative assistance. We also thank Drs. Karine Guillem, Magalie Lenoir and Youna Vandaele for their helpful comments on a previous version of the manuscript.

## Funding and Disclosure

The authors declare that they have no conflict of interest. This work was supported by the French Research Council (CNRS), the Université de Bordeaux, the Ministère de l’Enseignement Supérieur et de la Recherche (MESR) and the Fondation pour la Recherche Médicale (FRM DPA20140629788).

## Notes

### Competing Interest Statement

The authors have declared no competing interest.

